# Characterization of late structural maturation with a neuroanatomical marker that considers both cortical thickness and intracortical myelination

**DOI:** 10.1101/2021.02.24.432645

**Authors:** Sophie Maingault, Antonietta Pepe, Bernard Mazoyer, Nathalie Tzourio-Mazoyer, Fabrice Crivello

**Affiliations:** Groupe d’Imagerie Neurofonctionnelle, Institut des Maladies Neurodégénératives, UMR5293, Université de Bordeaux, Bordeaux, France; Groupe d’Imagerie Neurofonctionnelle, Institut des Maladies Neurodégénératives, UMR5293, CNRS, Bordeaux, France; Groupe d’Imagerie Neurofonctionnelle, Institut des Maladies Neurodégénératives, UMR5293, CEA, Bordeaux, France; Centre Hospitalier Universitaire Pellegrin, Bordeaux, France

**Keywords:** MRI, brain anatomy, maturation, cortical thickness, intracortical myelination, early adulthood

## Abstract

The cortical ribbon changes throughout a person’s lifespan, with the most significant changes occurring during crucial development and aging periods. Changes during adulthood are rarely investigated due to the scarcity of neuroimaging data during this period. After childhood, the brain loses gray matter, which is evidenced by an apparent reduction in cortical thickness (CT); one factor of this thinning process is intense ongoing intracortical myelination (MYEL). Here, we report age-related changes in CT, MYEL, and their ratio in 447 participants aged 18 to 57 years (BIL&GIN cohort). We propose the CT/MYEL ratio to be a multimodal cortical maturation index (MATUR) capable of reflecting 1) stages during which CT and MYEL patterns diverge and 2) the regional differences in cortical maturation that occur in adulthood. Age mainly decreased CT in all cortical regions, with larger reductions occurring in the bilateral insular lobes, temporal and frontal poles, and cingulate cortices. Age led to a linear increase in MYEL in the entire cortex and larger increases in the primary motor, auditory, and visual cortices. The effects of age on the MATUR index were characterized by both linear and quadratic components. The linear component mimicked the pattern found in CT, with 1) a robust amplification of the global and regional effects of age on CT and 2) evidence of new bilateral linear decreases in the frontal and cortical cortices. Most importantly, age exhibited additional large quadratic effects on the MATUR index in the bilateral frontal (more prominent in the right hemisphere), parietal, temporal, and cingulate regions that were not highlighted by the CT metric. Thus, the MATUR index was more sensitive to age-related cortical structural changes during adulthood than was either CT or MYEL alone. As evidenced by the large quadratic component of the effect of age, the newly proposed maturation index dramatically improved the characterization of the regional cortical territories, uncovering the latest brain maturation steps that occur before stabilization and deterioration occur in mid- and late adulthood.

## 1. INTRODUCTION

It is currently well known that the cortical ribbon changes enormously throughout a person’s lifespan. Many studies have been conducted to identify the genetic and environmental factors underlying its macrostructural changes, with the most significant changes occurring during two crucial periods, namely, the development period, including childhood and adolescence, and the later aging period, primarily referring to senescence (Giedd et al., 1999) (Sowell et al., 1999) (Walhovd et al., 2016). Between these two critical periods of life, the cortical ribbon changes that occur during adulthood, and more specifically during early adulthood, have rarely been investigated due to the relative scarcity of neuroimaging data during this period. However, the dramatic changes in the brain cortical mantle that occur during early development, both at the global and regional levels (Walhovd et al., 2016), continue throughout the postadolescence period (Norbom et al., 2020). This brain maturation stage involves fine-tuning structural changes and reorganization, which is why adults are more cognitively capable than are adolescents. Indeed, an adult brain differs from an adolescent brain in many ways. Between childhood and adulthood, the brain loses gray matter, as excess neurons and synapses are pruned away. This gray matter loss is characterized by an apparent reduction in cortical thickness (CT). The decrease in CT results from the combination of neuronal loss, synaptic pruning with a reduction in glial cells, and myelin proliferation into the cortical neuropil, with an increase in the caliber of the existing sheaths (Tamnes et al., 2010a) (Tamnes et al., 2010b). During early adulthood, the regional dynamics of the thinning of the cortical ribbon are nonlinear, being especially prominent in the parietal lobes and the medial and superior parts of the frontal lobe (Sowell et al., 1999) (Sowell et al., 2003) (Tamnes et al., 2010b). Afterward, thinning decelerates from when people are in their late 20s to when are in their early 40s (Westlye et al., 2010) (Fjell et al., 2013). In a modeling study, the effects of age on CT were represented by an exponential function, and the results demonstrated that 90% of thinning occurs before the age of 30 years in a broad set of regions, including the cuneus, the inferior temporal region, lateral occipital region, lateral orbitofrontal region, orbital part of the inferior frontal gyrus, and pericalcarine cortices, as well as the postcentral area and the superior parietal and frontal lobes (Tamnes et al., 2010b).

Early adulthood is thus a period of transition. The combination of advantageous maturational thinning and aging atrophy leads to a deleterious loss of neurons or cell body shrinkage and a reduction in synaptic density and the loss of myelinated axons (Morrison & Hof, 1997) (Hof & Morrison, 2004) (Lemaitre et al., 2012) (Hogstrom et al., 2013). In other words, the apparent thinning of the cortical ribbon estimated by MRI during adulthood is due to both late brain maturation and early aging atrophy processes; dissociating these two components is impossible when only CT measurements are considered.

Different neurobiological phenomena affect CT, and one factor contributing to CT thinning during early adulthood is the intense intracortical myelination process that occurs during this period. Intracortical myelination has been shown to reflect fiber density and caliber (Dinse et al., 2015). This intracortical myelination process is more pronounced in deep cortical layers close to the gray-white boundary, suggesting that it may contribute to the apparent thinning of the cortical ribbon associated with maturation (Natu et al., 2018). T1-weighted myelin-related magnetic resonance contrast images can reveal differences in lipids, free and myelin-bound water, and iron content colocalized with myelin within the cortical gray matter (Fukunaga et al., 2010). This method of imaging intracortical myelination has been validated using animal models as well as in human postmortem investigations, which have shown that there is a positive correlation between T1-weighted intensities and myelin content measured in brain scan slices (Bock et al., 2013) (Geyer et al., 2011) (Eickhoff et al., 2005). Such a correlation is higher in the internal half of the cortical ribbon, where axons emerge from the cortical ribbon (Eickhoff et al., 2005). This 3-stage profile has also been described by Grydeland et al. (Grydeland et al., 2013). This finding is also supported by the recent reports by Shafee et al. of an age-related linear increase in intracortical myelination in a population of 1,555 young adults aged 18 to 35 years (Shafee et al., 2015) and by Rowley et al. (Rowley et al., 2017), who showed that the intracortical myelin signal intensity peaks around the third decade but that the exact time depends on the region. Because the age-related trajectories of CT and intracortical myelination during adulthood differ, investigations on both of these two cortical phenotypes need to be conducted, as information on both factors could help elucidate the respective roles of maturational cortical thinning and aging-related atrophy during early adulthood and midlife.

In the present work, we investigated the relationship between CT and intracortical myelination as well as their joint age-related changes in a cohort of 447 healthy participants aged 18 to 57 years. We also investigate the age-related changes of the CT and intracortical myelination ratio, this latter index was considered a multimodal cortical maturation index capable of revealing periods during which CT and intracortical myelin patterns diverge.

These relationships were studied at the whole cortex level as well as at the cortical vertex level to determine the regional pattern of cortical maturation in adulthood.

## 2. MATERIALS AND METHODS

### 2.1. Participants

We conducted the present study using data collected from 447 participants and recorded in the BIL&GIN (http://www.gin.cnrs.fr/BILandGIN), a multimodal database designed for the study of human brain lateralization (Mazoyer et al., 2016). We recruited participants aged 18 to 57 years (mean = 26.7 ± 7.7 years) through announcements at Caen University. At the start of the study, none of the participants had neuropsychiatric disorders, were taking medications, or had structural brain MRI abnormalities. We conducted the study following the French laws on ethics in biomedical research: Basse-Normandie’s ethics committee approved the protocol. All participants gave their informed written consent and were compensated for their participation. The cohort was balanced in terms of handedness and sex (among the 243 right-handers, 114 were men, and among the 204 left-handers, 104 were men). The participants had 15.1 ± 2.5 years of education, which corresponds to 3 years of university-level education.

### 2.2. Image acquisition

The imaging scans were performed with a Philips Achieva 3 Tesla MRI scanner. The structural MRI protocol consisted of a localizer scan, high-resolution 3D-FFE-TFE T1-weighted volume acquisition (TR = 20 ms; TE = 4.6 ms; flip angle = 10°; inversion time = 800 ms; turbo field echo factor = 65; sense factor = 2; matrix size = 256 x 256 x 180; 1 mm^3^ isotropic voxel size) and 2D-TSE T2-weighted volume acquisition (TR = 5,500 ms; TE = 80 ms; sense factor = 2; FOV = 256 mm; matrix size = 256 x 256 x 81; 1 x 1 x 2 mm^3^ voxel size).

### 2.3. Image processing

#### 2.3.1. Surface-based processing

White matter and pial surface reconstruction were performed for each participant using the “recon-all” command in FreeSurfer 5.3.0 and the T1-weighted volume (http://surfer.nmr.mgh.harvard.edu/, (Dale et al., 1999) (Fischl et al., 1999) (Fischl & Dale, 2000). The FreeSurfer analysis included intensity bias field removal to optimize the present 3T acquisitions (−3T flag in the recon-all command line).

#### 2.3.2. Quality control

The surfaces around the thin temporal lobe strands that border the lateral ventricle tip has been shown to be difficult to estimate with the FreeSurfer segmentation procedure (Dale et al., 1999). A reduction in voxel intensity in the corresponding white matter leads to the misclassification of voxels and inaccurate identification of cortical surfaces. To overcome this limitation, one investigator (SM) visually checked each of the 447 pial and white surfaces on each axial, sagittal, and coronal section of each participant twice (Maingault et al., 2016). We identified segmentation errors in 48 participants (11%). We manually corrected the errors by adding an average of 9 +/− 7 control points to the misclassified white matter in all these cases. The automatic “recon-all” FreeSurfer procedure was rerun on these corrected volumes, resulting in a complete set of accurate surface reconstructions after the visual quality control step.

We recorded the total intracranial volume as a direct output of the FreeSurfer pipeline (mean = 1413 +− 141 cm^3^).

#### 2.3.3. Generation of the surface-based cortical thickness (CT) maps

For each individual, we generated a CT surface map. The CT values were integrated over all vertices of the cortex surface to determine the mean CT. A cortical surface curvature (CURV) map was also computed and used for nonlinear surface-based between-individual registration.

The 447 CT individual maps were then spatially normalized to a custom surface-based template specifically based on the MRI data of 40 participants included in the BIL&GIN. These participants were representative of the entire study population in terms of age, sex, and handedness (for a detailed description of the specific surface-based template, see Marie et al. (Marie et al., 2016)).

#### 2.3.4. Generation of the surface-based intracortical myelin (MYEL) maps

Each T2-weighted (T2w) volume was rigidly registered to the corresponding T1-weighted (T1w) volume using the “SPM12 coregister” routine and resampled (isotropic voxel size of 1 mm^3^) using trilinear interpolation (http://www.fil.ion.ucl.ac.uk/spm/). For each participant, a T1w/T2w ratio volume was computed and sampled onto the subject cortical surface by taking the T1w/T2w value at a mid-thickness distance into the gray matter ribbon (Glasser & Van Essen, 2011) (Grydeland et al., 2013), yielding a T1w/T2w ratio surface map in the individual participant’s space. We performed this sampling technique using the same surface reconstruction method as that used for generating the CT maps, thus ensuring perfect vertex correspondence between CT and T1w/T2w ratio surface maps. We assigned a T1w/T2w value of zero to vertices located in the noncortical medial wall or those that had a null CT value. The T1w/T2w surface map intensities were normalized across subjects using the mean value of the T1w/T2w ratio in the deep white matter region, as defined by the FreeSurfer white matter parcellation step. Hereafter, the intensity-normalized T1w/T2w surface maps will be referred to as the MYEL maps.

We calculated the global cortex MYEL value by averaging the vertexwise nonnull MYEL values over the entire cortex. Finally, we spatially normalized all 447 individual MYEL surface maps to the BIL&GIN-specific surface-based template. The resulting 447 MYEL maps represented a proxy for the intracortical myelin content of the BIL&GIN participants. Note that MYEL is a unitless quantity defined as the ratio of unitless raw intensity data.

#### 2.3.5. Definition of the surface-based multimodal cortical maturation index (MATUR) maps

For each participant, we defined a vertexwise multimodal cortical maturation index (MATUR) map by computing the ratio between the individual CT and MYEL (in mm) vertexwise maps, excluding the vertices with null values for MYEL. For CT and MYEL, we resampled the 447 MATUR maps with the BIL&GIN-specific surface-based template.

### 2.4. Statistical analysis

#### 2.4.1. Descriptive statistics

We computed the average values of the CT, MYEL, and MATUR variables at the whole cortex level and the vertex level across all participants, resulting in average surface maps.

In addition, we specifically examined the regional relationship between CT and MYEL using linear regression modeling. We assessed the correlations between MYEL and CT after the smoothing of the individual CT and MYEL maps with a small 2 mm full-width-at-half-maximum (FWHM) Gaussian kernel (to preserve the best spatial resolution available). We performed these computations using the MATLAB “partialcorr” correlation function with the vertexwise CURV values as the per-vertex regressor and age as the global regressor. A false discovery rate (FDR) threshold was set at q=0.05 and corrected for multiple comparisons to highlight the loci exhibiting significant correlations at the vertex level.

#### 2.4.2. Age effects

The effects of age on each of the three variables, CT, MYEL, and MATUR, were assessed both at the global level (*i*.*e*., using the average value of the variable over the whole cortex) and at the vertex level (*i*.*e*., measuring the effects of age at each vertex of the surface map). At the whole cortex level and in all cases, we assessed the effects of age using regression analysis and by comparing two models, one including a simple linear age effect and the other including an additional age-squared effect in addition to the linear age effect. In each case, the model exhibiting the lowest corrected Akaike information criterion (AICc, Akaike, 1974) was retained, and the sizes of the age and age-squared effects were measured using the partial eta-squared statistic (Lakens, 2013). The age effects on the whole cortex at the 90% level were determined using JMP® software (version *Pro13*.*0*. SAS Institute Inc., Cary, NC, 1989-2019), and the size of the effects were determined with the “*Calculate Effects Sizes*” JMP add in (version 0.07, https://community.jmp.com/t5/JMP-Add-Ins/Calculate-Effect-Sizes-Add-in/ta-p/22642). Ninety percent confidence intervals of the effect sizes were computed with the “MBESS” R package (Ken, 2007). The 90% confidence interval was equivalent to the 95% two-sided confidence interval, as the F-statistic cannot be negative (Smithson, 2003).

To assess effects of age at the vertex level, the CT, MYEL, and MATUR surface maps were first smoothed with a 15 mm FWHM Gaussian kernel. Linear modeling was then performed at each vertex site using the “*mri_glmfit*” command in the FreeSurfer package to test the linear and quadratic effects of age on the CT, MYEL, and MATUR variables. Both predictor variables (age and age^2^) were entered into an ANOVA model.

The statistical maps of each predictor were thresholded at q=0.05 (FDR correction for multiple comparisons) to identify the significant linear and quadratic effects of age on CT, MYEL, and MATUR. These “q-value corrected maps” were then used to mask the vertexwise effect maps (*i*.*e*., the regression coefficients of both age and age^2^ effect).

## 3. RESULTS

### 3.1. Descriptive statistics

#### 3.1.1. Cortical thickness (CT)

The average mean CT was 2.44 ± 0.11 mm (mean ± standard deviation). The average CT surface map (N=447) of the study population is shown in Figure 1 (A). We found that the cortex was thicker in the precentral gyrus, inferior frontal gyrus, and dorsal part of the medial surface of the frontal lobes. In the parietal lobe, the cortex was thicker in the postcentral sulcus and the supramarginal and angular gyri, as well as internally in the posterior part of the precuneus and in the posterior part of the cingulate gyrus.

**Figure 1.**
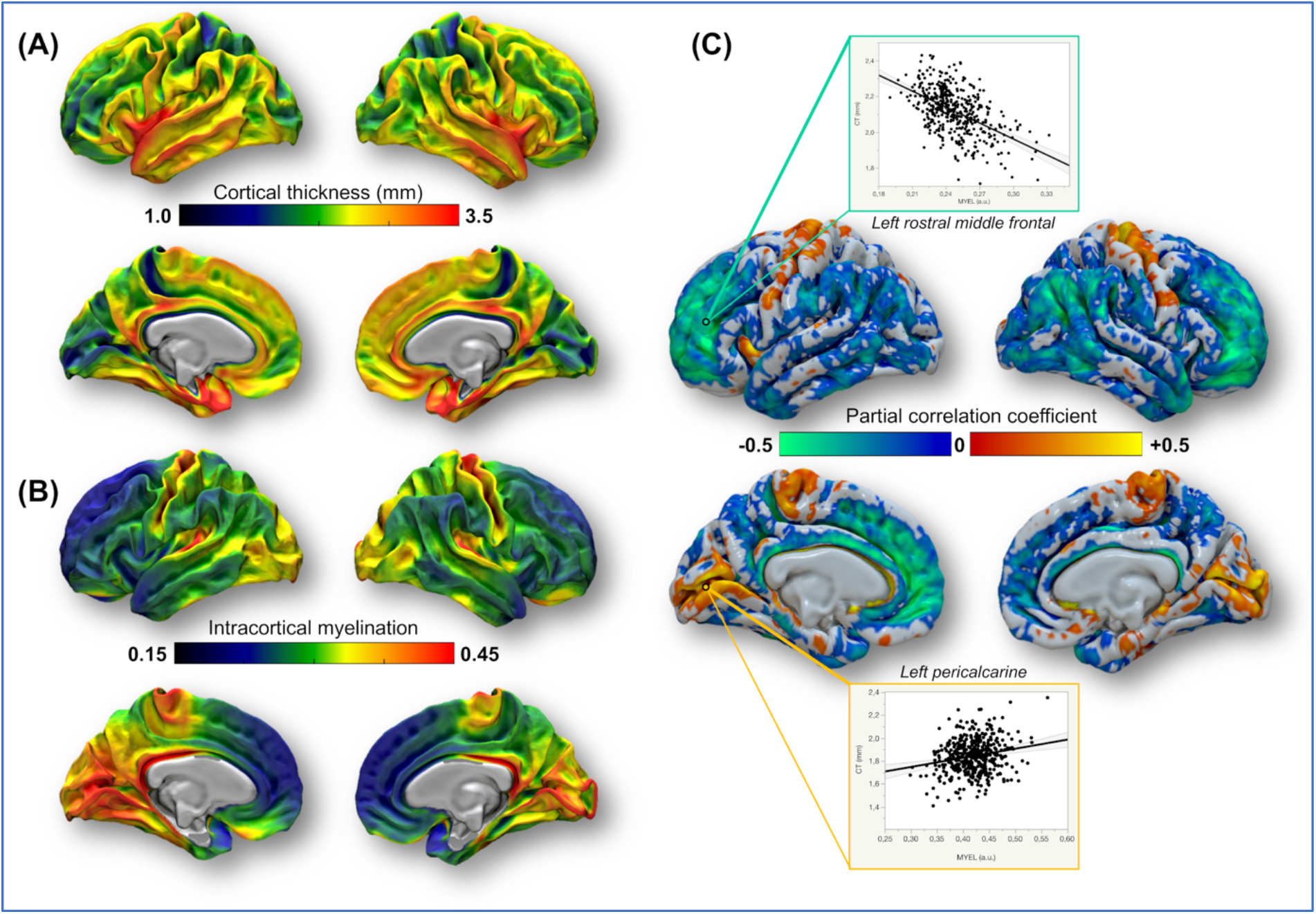
**A)** and **B)**. Average maps of cortical thickness (CT), expressed in mm, and intracortical myelin content (MYEL) projected on the reconstructed white surfaces of both hemispheres in the surface-based reference space averaged across 40 participants. **C)**. Correlation map of CT and MYEL controlled for curvature. Only the partial correlation coefficients meeting the FDR threshold of q = 0.05 are presented. Both the positive (hot colors) and negative (cold colors) correlations are displayed and projected on the reconstructed pial surfaces of both hemispheres of the surface-based reference space averaged across 40 participants.

#### 3.1.2. Intracortical myelin

The average mean intracortical myelin value was 0.28 ± 0.03 a.u. (mean ± standard deviation). The average MYEL surface map for the study population is shown in Figure 1 (B part). Regions with high intracortical myelination values were located bilaterally in the cortices of the primary areas, including the motor area along the rolandic sulcus, the visual supplementary motor area along the calcarine fissure and towards the temporal pole in the lingual and fusiform gyri, and auditory supplementary motor area around Heschl’s gyri. The temporal lobe showed high intracortical myelination on its lateral and medial surfaces, with peaks on the temporal poles. Finally, the insular lobe also presented high values of intracortical myelination. In the medial surface of the hemispheres, the posterior half of the cingulate gyrus also showed high myelination bilaterally.

#### 3.1.3. Correlation between cortical thickness (CT) and intracortical myelin

At the whole cortex level, CT and MYEL were negatively correlated (R^2^ = 0.31, p <0.0001). We found that a very large part of the cortex was characterized by strong negative correlations between the vertexwise CT and MYEL maps, indicating that the thicker the cortex was, the smaller the intracortical myelin content (see Figure 1, part C). The regions showing the strongest negative correlations were located bilaterally in the middle and superior frontal gyrus, the supramarginal and inferior parietal gyrus, and the inferior and middle temporal gyrus. On the internal surface, strong negative correlations were also found in the anterior cingulate cortex (mainly on the left hemisphere) and bilaterally in the precuneus. Only a small set of regions exhibited positive correlations between CT and MYEL. These regions surrounded the primary cortical areas, notably including the pericalcarine cortex and the pre-and postcentral gyri.

#### 3.1.4 Maturation index

The average mean maturation index was 8.21 ± 0.92 (mean ± standard deviation). The average MATUR surface map is shown in Figure 2. The highest MATUR values were located in the superior frontal gyrus, particularly in its medial part; the cingulate cortex; the precentral gyrus; the supramarginal and angular gyri; the anterior part of the temporal lobe; along the middle temporal gyrus; and the insula.

**Figure 2.**
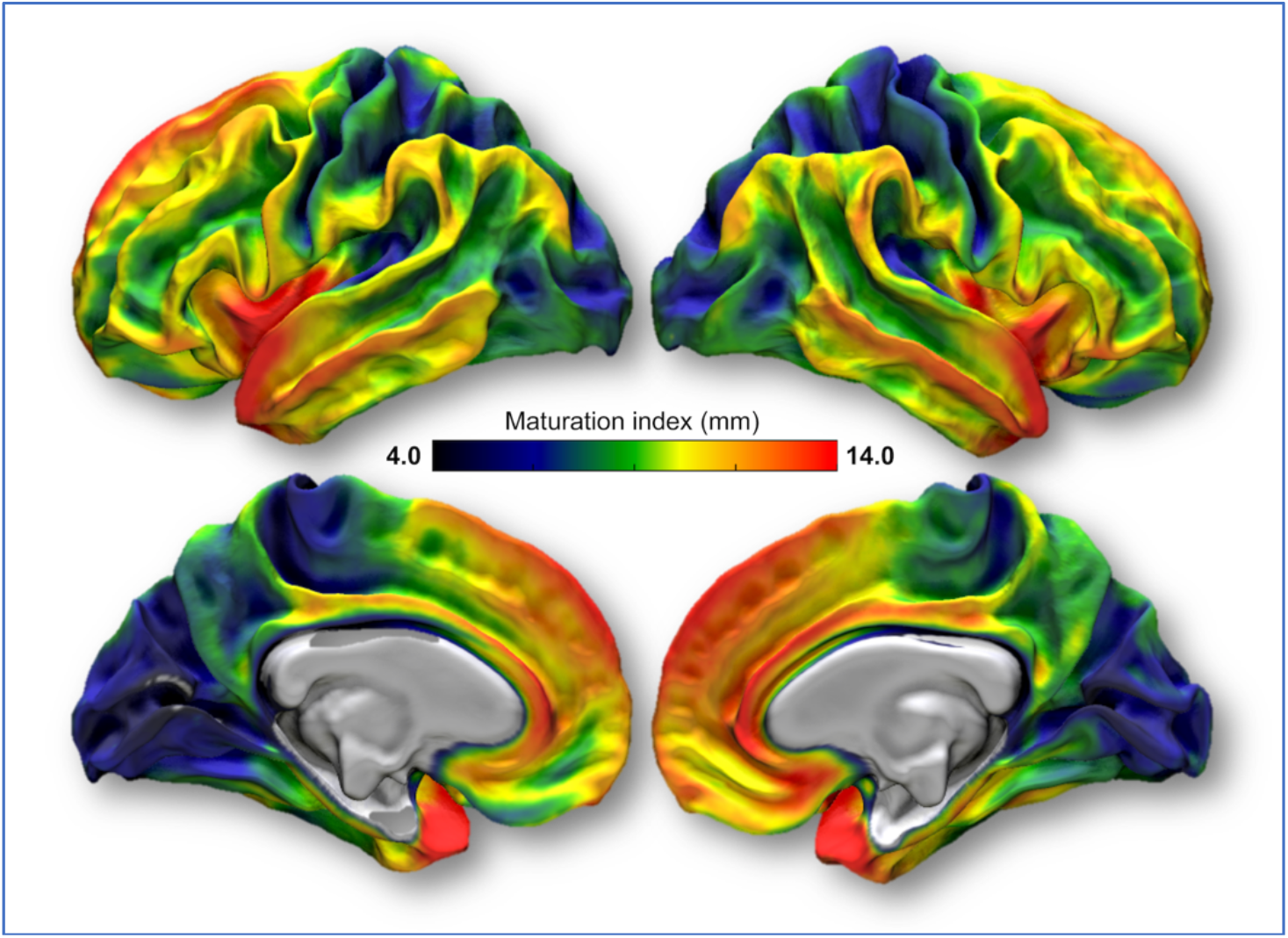
The average map of the newly proposed cortical maturation index (MATUR), expressed in mm, projected on the reconstructed white surfaces of both hemispheres of the surface-based reference space averaged across 40 participants.

In contrast, the lowest values of this index were observed in the whole medial part of the occipital lobe containing the pericalcarine cortex, the area of the central sulcus extending to the marginal sulcus, the transverse temporal gyrus, and the intraparietal sulcus.

### 3.2. Age effects

#### 3.2.1. Age effect on cortical thickness (CT)

The mean CT was found to be best fitted by a quadratic model, as assessed by the AICc (see Table 1). The quadratic relationship between CT and age (R^2^ = 0.30, p < 0.0001) is illustrated in Figure 3 (panel A). The mean decrease in CT was thus characterized by both significant linear age and age^2^ effects (β_age_ = −9.6 μm.year^−1^, p < 0.0001 and β_age2_ = +0.16 μm.year^−2^, p = 0.0029). The positive value for the age^2^ effect indicates a deceleration in CT thinning between 18 and 57 years; in other words, there was a larger decrease per year in the youngest subjects than in the older subjects. However, the age^2^ effect size was small (approximately 2% of explained variance) compared to the linear age effect size, which was approximately ten times larger (see Table 1).

**Table 1.** Size of the age and age^2^ effects at the whole cortex level

| | AICc | | $\eta^2$ | |
| --- | --- | --- | --- | --- |
|  | linear | quadratic | age | age<sup>2</sup> |
| CT | -870.58 | **-877.46** | 0.20 [0.15, 0.25] | 0.02 [0.00, 0.05] |
| MYEL | **-2273.08** | -2273.00 | 0.25 [0.20, 0.301] |  |
| MATUR | +947.17 | **+936.97** | 0.31 [0.25, 0.36] | 0.03 [0.01, 0.06] |

CT: cortical thickness; MYEL: intracortical myelin; MATUR: maturation index.
AICc: Akaike information criterion; the best model is in bold font.
linear: variable = intercept + age + error.
quadratic: variable = intercept + age + age<sup>2</sup> + error.
$\eta^2$ : partial eta-squared statistic and 90% confidence interval.

**Figure 3.**
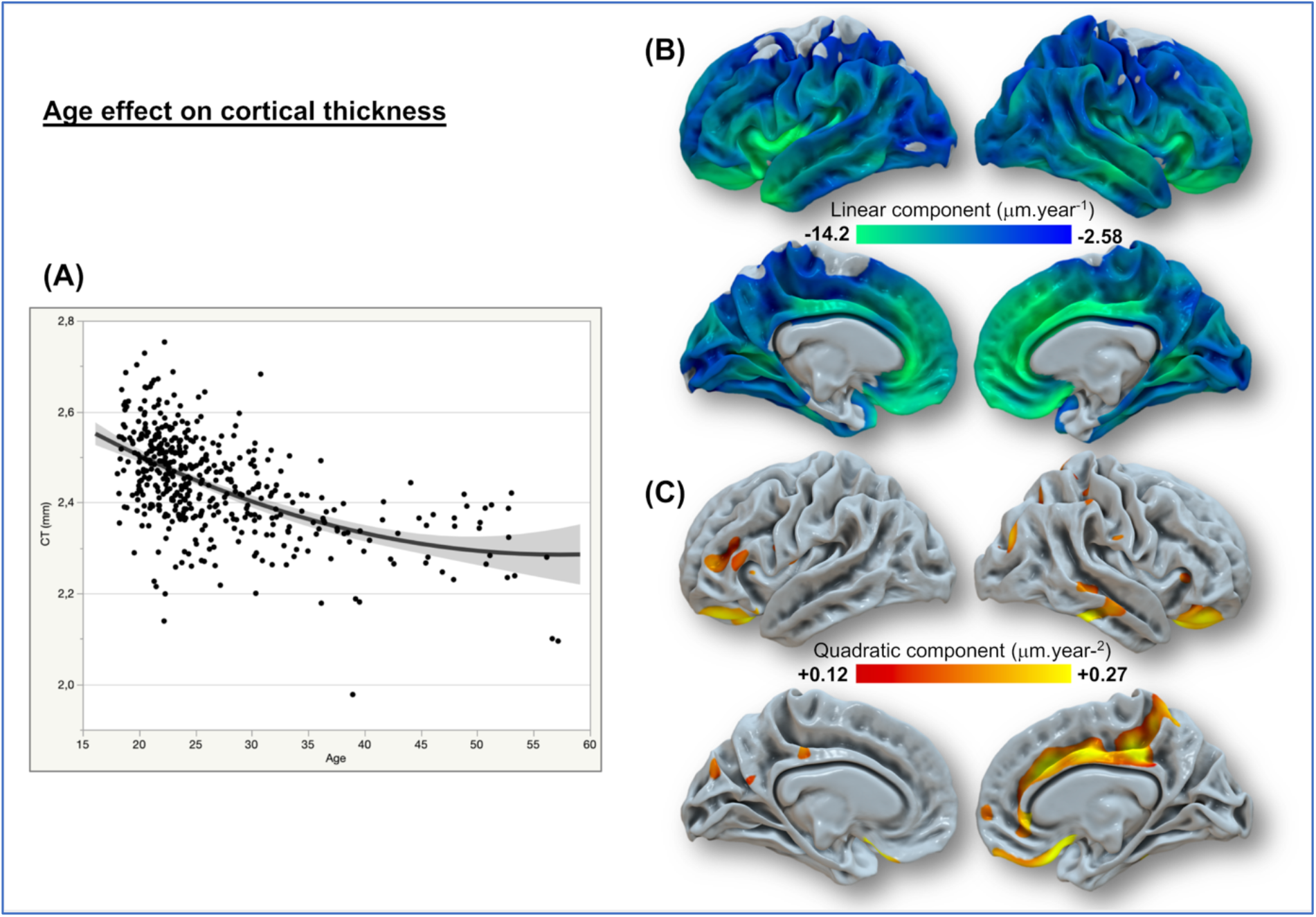
Age-related cortical thickness (CT) changes. **A)** Quadratic fit of the mean CT decreasing with age. **B)** Vertexwise spatial distribution of CT linear thinning with age (expressed in mm per year). **C)** Vertexwise spatial distribution of the CT-positive quadratic age effect (defined in mm per year^2^). Only the age effects meeting the FDR threshold of q = 0.05 are presented and projected on the reconstructed white surfaces of both hemispheres of the surface-based reference space averaged across 40 participants.

At the vertex level, the effect of age was mainly characterized by a linear effect with a relatively symmetric cortical thinning pattern (Figure 3, panel B). Note that no regions exhibited an increase in CT with age. The linear negative association between age and CT was observed in almost all cortical areas, with the highest rate of thinning being observed in the insula and the frontal lobe area extending along the cingulate gyri, including the medial orbital frontal and superior regions, the inferior frontal gyrus, and the rostral part of the middle frontal gyri. In the parietal lobe, the supramarginal gyrus, the cuneus, and the precuneus showed high thinning CT rates, whereas the CT in the postcentral region appeared to be almost stable. Higher rates of CT thinning were also found in the temporal pole and at the level of the superior temporal sulcus (more pronounced in the right hemisphere). Finally, the occipital regions presented relatively stable CT values.

Significant positive quadratic effects of age on CT were observed in few small clusters (no significant negative age^2^ effect clusters were detected). These clusters were located in the left hemisphere in the rostral part of the middle frontal gyrus, the pars triangularis of the inferior frontal gyrus, the orbital gyrus, the precuneus and the posterior cingulate gyrus. In the right hemisphere, the clusters were located in the anterior, middle and posterior cingulate gyrus, the paracentral lobule, the orbital gyrus and in the mid part of the right middle temporal gyrus (Figure 3, panel C). Such sparse and weak age^2^ effects at the voxel level explain the weak age^2^ effect at the whole cortex level.

#### 3.2.2. Age effects on intracortical myelin

Contrary to the mean CT, the mean MYEL increased with age and was best fitted by a linear model (see Table 1). The linear relationship of MYEL with age (R^2^ = 0.44, p < 0.0001) is illustrated in Figure 4 (panel A), and the effect of age on MYEL was significant (β_age_ = 21.31×10^−^4/year, p < 0.0001), while the age^2^ effect in the quadratic model did not reach significance (p = 0.16). Age explained 25% of the variance in MYEL, which was higher than the percent of variance in CT explained by age (see Table 1).

**Figure 4.**
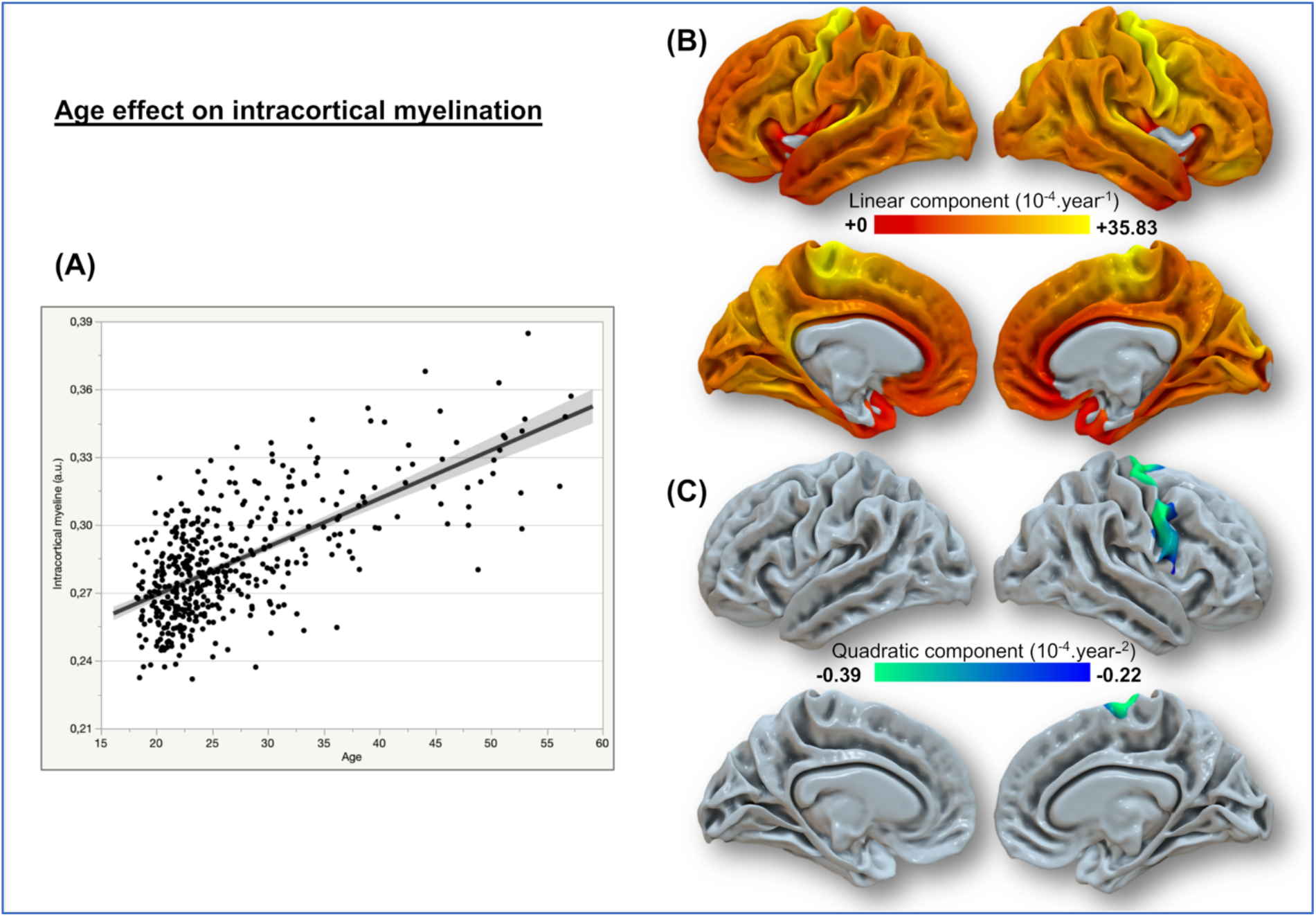
Age-related intracortical myelin (MYEL) changes. **A)** Linear fit of the mean MYEL increasing with age. **B)** Vertexwise spatial distribution of MYEL linearly increasing with age (defined per year). **C)** Vertexwise spatial distribution of the MYEL negative quadratic age effect (defined per year^2^). Only the age effects meeting the FDR threshold of q = 0.05 are presented and projected on the reconstructed white surfaces of both hemispheres of the surface-based reference space averaged across 40 participants.

At the vertex level, almost the whole cortex showed a significant linear relationship with age; MYEL increased with age, while CT decreased with age. The regions with the highest growth rate were located bilaterally in the precentral gyrus (primary motor cortex) area extending to the paracentral lobule in the medial part of the hemisphere and in the transverse temporal (Heschl’s) gyrus in the primary auditory cortex. Conversely, the anterior part of the cingulate gyrus, the gyrus rectus, and the temporal pole displayed the smallest increases that were still considered significant. Note that the absence of a significant effect in the bilateral insula was due to the statistical threshold used for assessing the effect of age, and the insula showed the smallest linear increase in MYEL (Figure 4, panel B).

Only two brain areas exhibited a significant age^2^ effect on MYEL: the right inferior part of the precentral gyrus and a region extending internally to the right supplementary motor area (Figure 4, panel C). The negative values of the age^2^ effect in these specific regions indicate a smaller MYEL increase in the oldest participants than in the youngest participants. The fact that significant age^2^ effects were observed only in this restricted set of areas explains why there was no age^2^ effect on MYEL at the whole cortex level.

#### 3.2.3. Age effect on the maturation index

For CT, the whole cortex mean MATUR index was found to be best fitted using a quadratic model (see Table 1). The mean MATUR decrease was thus characterized by both significant age and age^2^ predictors (β_age_ = −97.1 μm.year^−1^, p <0.0001 and β_age2_ = +1.4 μm.year^−2^, p = 0.0005, respectively), revealing a large decrease in the MATUR index between 18 and 57 years, and also a deceleration of this decrease.

Across the whole cortex, MATUR showed a significant negative linear relationship with age (Figure 5, panel A). The goodness of fit value for the quadratic model was significantly larger for the mean MATUR index for the whole cortex than for the mean CT (R^2^ = 0.44 *vs*. 0.30 for MATUR and CT, respectively). The effect size of age on MATUR was +53.4% higher than that on the mean CT (see Table 1). The age^2^ effect size was also larger (+36.7%) on MATUR than on CT.

**Figure 5.**
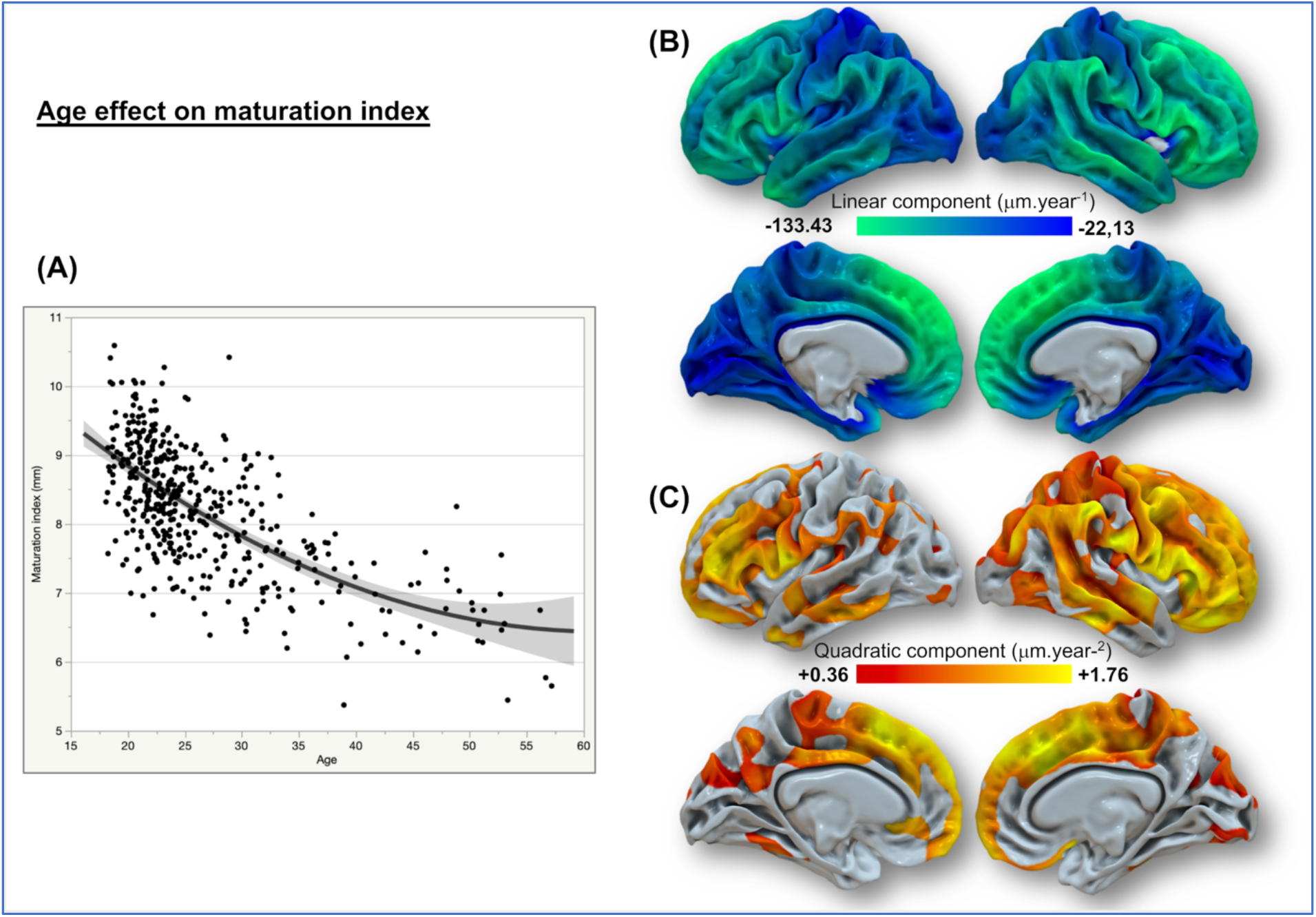
Age-related cortical maturation index (MATUR) changes. **A)** Quadratic fit of the mean MATUR decreasing with age. **B)** Vertexwise spatial distribution of MATUR linear thinning with age (expressed in mm per year). **C)** Vertexwise spatial distribution of the MATUR-positive quadratic age effect (defined in mm per year^2^). Only the age effects meeting the FDR threshold of q = 0.05 are presented and projected on the reconstructed white surfaces of both hemispheres of the surface-based reference space averaged across 40 participants. The positive quadratic age effect on MATUR depicts the territories where the younger participants presented a significantly larger MATUR decrease than did the older participants.

The vertex maps of the linear effect of age on MATUR revealed regional heterogeneity (Figure 5, panel B). The highest rates of linear decline were observed in the whole frontal lobe, including the cingulate regions, with a relative rightward predominance. The parietal lobe; the parietal operculum; the inferior parietal gyrus, including the supramarginal gyrus; and the angular gyrus were the sites of the largest rates of linear decline. In the temporal lobe, large linear decreases with age were also found in the right middle and inferior temporal gyri and in the left temporal pole and posterior part of the inferior temporal gyrus.

Contrary to the age^2^ effect on CT, that on MATUR was significant and positive in large parts of the cortex, suggesting that MATUR decreased more in the youngest subjects than in the older subjects in extensive areas (Figure 5, panel C). This decline in the MATUR rate of decrease was observed in the frontal lobes; in the present study, a clear rightward predominance centered on the middle and inferior frontal gyri, the rolandic operculum, and medially in the superior frontal gyrus around the lateral part of the gyrus was observed. In the right parietal lobe, the parietal operculum; the inferior parietal gyrus; including the supra-marginal gyrus and the angular gyrus; and internally, the precuneus had the highest rates of decrease. In the temporal lobe, the highest quadratic effects of age were observed in the posterior two-thirds of the right middle temporal gyrus and in the left in the temporal pole. Note that the cortical areas showing the largest quadratic effect of age on MATUR also showed the largest negative linear effect of age. Similarly, the brain areas displaying no significant age^2^ effect, essentially the occipital lobes, also exhibited the smallest linear decrease in MATUR.

### 3.3. Summary of the results

In almost all regions (especially in the frontal, parietal and cingulate cortices), CT decreased during early and mid-adulthood. Moreover, myelination increased, except in the visual and motor primary cortices and supplementary motor areas where the CT and MYEL increased. Consequently, the MATUR index, which reflects the ratio of these parameters, presented substantial regional variation, with its highest values being observed 1) in the cortices mentioned above, showing a CT/MYEL negative association, and 2) in the regions with the highest CT values (insula lobe and temporal cortices). In contrast, the lowest values of the MATUR index were observed in 1) the cortices showing a positive association between CT and MYEL and 2) regions presenting the highest CT values (primary motor, visual, and auditory cortices). The CT mostly decreased linearly in all cortical regions with age, with a larger decrease in the bilateral insular lobes, temporal and frontal poles, and cingulate cortices. MYEL linearly decreased with age in the entire cortex, with a larger increase in the primary motor, auditory, and visual cortices. The effects of age on the MATUR index were characterized by both linear and quadratic components. The pattern of the linear component mimicked that of the effect of age on CT, with 1) a strong amplification of the regional effect of age on CT and 2) evidence of a new large linear decrease in the bilateral frontal and cortical cortices. Most importantly, age exhibited additional large quadratic effects on the MATUR index in the bilateral frontal, parietal, temporal, and cingular regions, which were not reflected by the CT metric.

## 4. DISCUSSION

### 4.1. Characterization of cortical thickness (CT) and intracortical myelination patterns, methodological issues, and limitations of the study

The mean CT pattern observed in the present study is in line with the pioneer map described by Fischl et al., which showed thick insula and temporal lobe areas and thin frontal, somatosensory (S1), and visual (V1) areas (Fischl & Dale, 2000). The current CT map also allows the thicker somatosensory (S1) area to be differentiated from the thicker motor (M1) areas by manual measurements of CT from MRI data (Meyer et al., 1996). Regarding the mean CT pattern’s spatial characteristics, we observed a thinner cortex at the depths of the sulci and a thicker cortex at the gyri crowns. These results are consistent with those of a postmortem study using high-field-strength MRI slice data of the brain (Fatterpekar et al., 2002). According to the tension-based theory of folding developed by Van Essen, cells in deep layers should be stretched tangentially and compressed radially along inward folds or sulci, making these layers thinner. In contrast, along the outer folds corresponding to gyri, the cells in deep layers should be stretched radially, making them thicker (Van Essen, 1997). The proposed CT pattern is in complete agreement with the one presented in Valk et al.’s study (Valk et al., 2017), who computed the mean CT in 392 healthy adults with nearly balanced proportions of males and females and an age rage (from 20 to 55 years) very similar to that in our study (from 18 to 57 years).

The T1w over T2w maps generated in the present study were used to estimate the mean local intracortical myelination value. In previous studies, Grydeland measured intracortical myelin at a distance of 0.2 mm above the gray matter/white matter interface (Grydeland et al., 2013) (Grydeland et al., 2016), but we estimated the surface-based representation of this measure by assessing the intracortical myelin signal at the area of mid-thickness in the cortical ribbon. The MYEL spatial distribution in our study showed that this methodological step performed to reduce the partial volume effect potentially introduced by 2 mm T2w slice thickness MRI acquisition did not alter the intracortical myelin representation content.

Indeed, we observed lower myelin content in both unimodal and heteromodal associative areas located in the temporal, parietal, and frontal lobes coupled with greater myelination in primary cortical areas. This pattern is consistent with the myelin content patterns presented in previous studies (Glasser & Van Essen, 2011) (Grydeland et al., 2013) (Ganzetti et al., 2014) (Shafee et al., 2015) (Norbom et al., 2020). However, we cannot exclude the fact that the lower spatial resolution of the T2w images in the present study may have caused regional partial volume effects, especially in thin and heavily myelinated areas such as the pericalcarine cortex. Indeed, our results showed a large area with a high MYEL signal covering the whole medial part of the occipital lobe (Figure 1, B part). De Martino et al. determined the mean T1w/T2w ratio on surface-based images (N=6) using 0.6 mm isotropic high-resolution acquisition with a 7T scanner (De Martino et al., 2015). The authors reported a lower signal in the occipital lobe than in the central sulcus or the paracentral lobule, suggesting that the MYEL signal deteriorated in our study due to a specific partial volume effect in this extremely thin region. Moreover, based on high-resolution imaging, Rowley et al. proposed a method of separating the cortical ribbon along its depth to differentiate lightly and heavily myelinated portions (Rowley et al., 2015). The authors reported a restricted occipital territory compared to the one presented in our study that was explicitly located around the calcarine fissure, did not extend to the cuneus, and had a high myelinated thickness. Nevertheless, despite the relatively low spatial resolution of the T2w acquisition system used in the present study, the MYEL mean map presented a fair final spatial resolution, highlighting quite precisely the focal points of high intracortical myelination values concentrated in the primary cortices. Another illustration of the consistency of the current MYEL maps at the macroanatomical scale was provided in a previous article using the same database, in which different patterns of intracortical myelin content were associated with the variation in Heschl’s gyri duplication at the individual level (Tzourio-Mazoyer et al., 2018). In this study, the high MYEL signal in the auditory areas on the average map was consistent with the fine-grained description of intracortical myelination provided by Glasser et al. (Glasser & Van Essen, 2011).

Although the present MYEL map was carefully characterized, one limitation of the present study is that the resolution of the MRI scans acquired between 2007 and 2008 were relatively coarse compared with what can currently be achieved with a comparable acquisition time. In addition, absolute quantitative T1 acquisition systems are now available, which can better reveal intracortical myelination markers (Lutti et al., 2014) (Dinse et al., 2015) (Huntenburg et al., 2017).

Finally, we specifically created an adequate surface-based template for the group vertexwise analysis. It is widely acknowledged that individual images need to be normalized to a template created from data that are acquired with the same imaging system and have the same characteristics as the data to be analyzed for high study sensitivity and accuracy (Klein et al., 2009) (Klein et al., 2010). To exclude systematic template bias from the stereotaxic analyses, the template should be calculated from a population as similar as possible to the study population if not from the study population itself (Evans et al., 2012). The use of such a specific-population template reduces areal shifts and biases caused by registration (Winkler et al., 2012). The FreeSurfer template (namely, Fsaverage) was built from MRI data acquired from healthy individuals of ages ranging from 18 to 93 (and no subjects between 30 and 65 years of age) with an imaging system different from ours. To optimize our data analyses, we thus developed a specific template for our study population from a subset of individuals included in the BIL&GIN database balanced by sex and handedness.

### 4.2. Rationale for the use of a multimodal cortical maturation index based on both cortical thickness (CT) and intracortical myelination

According to Glasser et al. (Glasser & Van Essen, 2011), the T1w over T2w ratio can be considered a pertinent intracortical myelination marker. However, this marker appears to not be independent of other cortical ribbon structural features, including CT. Indeed, MYEL is strongly related to CT. In particular, Shafee et al. reported a vertexwise correlation between CT and MYEL after controlling for the local curvature (Shafee et al., 2015). The present study shows a strong negative correlation between MYEL and CT in almost all areas (principally in parietal, temporal, and frontal regions, see Figure 1 C part). The correlation pattern we observed agreed with the one proposed by Shaffe et al., revealing the same spatial correlation pattern. This feature has also been shown in a high-field high-resolution validation study combining cyto- and myeloarchitectonics with MRI (Dinse et al., 2015). Our MYEL/CT correlation analysis also highlights some specific positive correlations detected in the central sulcus and paracentral lobule, primary visual cortex, and parahippocampal cortices. This positive correlation was also observed by Shaffe et al., who attributed this positive correlation to the fact that these regions present the thinnest cortices. In these areas, the surface representing the GM/WM interface was mistakenly estimated, thus generating a thinner cortex and lower MYEL value and causing an artifactual positive correlation between CT and MYEL.

The negative correlation between MYEL and CT observed in our study was expected. Indeed, intracortical myelination plays a role in the cortical surface being stretched along the tangential axis, leading to thinning of the cortex (Lorio et al., 2016), which could explain the predominant negative correlation between CT and the intracortical myelin content. This phenomenon is hypothesized to disentangle neighboring neuronal columns, enable the afferent patterns (myelination) in the cortex to be differentiated well, and to consequently increase functional specialization. However, the link previously described between CT and MYEL seems to be more strongly related to the maturation process than to the developmental process observed during childhood. Using a myelin water fraction marker, Croteau-Chonka et al. investigated the dynamics of the relationship between CT and this white matter development proxy through early childhood (from 1 to 6 years) (Croteau-Chonka et al., 2016). These authors reported that in only 14 of 66 (21%) cortical regions analyzed in their study, changes in CT were correlated with changes in adjacent white matter. This negative correlation was weaker than that observed in our study, and this inconsistency could be due to 1) the low resolution of FreeSurfer’s parcellation, especially in children’s brains, with respect to vertexwise analysis; 2) the estimation of the myelin water fraction, which may be less sensitive to changes than the MYEL estimation; and 3) measures of CT and white matter maturation, which reflect distinct but complementary neurodevelopmental processes but do not characterize identical underlying processes.

We observed a stronger negative association between CT and MYEL in the bilateral superior and middle frontal regions, the parietal lobe, the lateral and posterior part of the temporal lobe, and the precuneus. These regions are the sites of heteromodal associative areas, which are the least well organized from a cytoarchitectonic point of view, unlike the primary cortices (Mesulam, 1998) and the regions that develop the latest during cerebral maturation (Amlien et al., 2015). These areas are also characterized by a recent expansion in human evolution and high postnatal cortical growth (Hill et al., 2010). Moreover, these regions present the highest sulcal depth variability, and this pattern is associated with the local intrinsic connectivity variability (Mueller et al., 2013). These factors suggest that associative heteromodal areas presenting the strongest CT and MYEL negative associations are subject to be impacted by postnatal environmental factors. This relationship may illustrate a determinant of the late maturation process in young adults.

Age-related cortical thinning is not just a deleterious phenomenon linked to brain atrophy and neuronal death. As previously described, cortical thinning is driven by increased myelination in the white matter tracts coursing within and near the deepest part of the cortical layer (Marsh et al., 2008) (Erus et al., 2015). This phenomenon occurs until 30 years of age and does not seem to progress during mid-adulthood (Grydeland et al., 2013). This process is thus clearly illustrated by the proposed maturation index, which considers both CT and MYEL. This CT to MYEL ratio amplifies the opposed temporal dynamics of CT and intracortical myelination (see Figures 3 and 4, respectively) and consequently, better characterizes the macrostructural changes during adulthood.

When considered separately, neither CT nor MYEL allowed us to sharply discriminate the age effects that may differ between young and middle-aged adults (Figures 3 and 4), which was mainly due to the relative scarcity of areas presenting a significant quadratic age effect.

However, the effects of age observed on CT or MYEL alone, which were mainly linear, are highly consistent with those reported in the literature. We found a considerable decrease in CT with age in high-order cortices, indicating that thinning occurs in these areas not only during adolescence but also during adulthood (Sowell et al., 1999). The CT thinning rates we observed were similar to those found in studies dealing with development and maturation, i.e., approximately 20 µm/year, which is two to three times higher than the rate of atrophy during senescence. Such an effect of age is consistent with the highest magnitudes of thinning in the heteromodal association cortices and regions of high postnatal surface area expansion in young adults (McGinnis et al., 2011). Overall, when considering the CT marker alone, we found a low regional specificity in terms of the different effects of age between young- and middle-aged adults, as revealed by the small extent of clusters showing significant quadratic effects of age.

Considering the MYEL metric, the age effect’s quadratic component was restricted to only one area, namely, the right precentral gyrus (Figure 4). This pattern corresponding to a deceleration in intracortical myelination increased with age in the precentral region, while almost all cortical areas showed a constant increase in myelination with age, as illustrated by the entire cortex coverage of the linear component of the age effect on MYEL. These results are consistent with those of a study recently conducted by Norbom et al. (Norbom et al., 2020). In that study, intracortical myelination changes in childhood and late adolescence (age from 3 to 21 years) were investigated, and the authors did not find any statistically significant associations between the T1w/T2w ratio and square of age when controlling for linear age effects. The present study also shows that the linear age effect is observed across the entire cortical mantle, both internally and laterally, except for in the medial occipital lobe. The linear component of the age effect on MYEL increased at a higher rate in both banks of the central sulcus, the paracentral lobule, and in the supratemporal plane bilaterally. These findings are also consistent with previous observations showing that the myelination of intracortical axons is high in primary cortical areas during childhood and adolescence (Norbom et al., 2020), during mid-adulthood (Rowley et al., 2017) and later in life, as it continues to increase until approximately 60 years of age (Grydeland et al., 2013).

### 4.3. Regional dynamics of maturation: significant changes in the high-order frontal and parietal regions in young adults

In contrast to the maps of CT changes, the MATUR index maps provided dramatically more sensitive estimations of cortical ribbon changes with age. These changes were illustrated by both linear and quadratic predictor components of the age effect and, therefore, allows better differentiation of the age effect between young adults and the oldest adults. First, we observed that the age effects (both linear and quadratic) on MATUR computed for the whole cortex were 50% larger than those on CT. Second, for the vertexwise linear age effect, we found a tenfold higher rate of decrease in MATUR (from −22 to −133 μm.year^−1^) than in CT (from −2 to −14 μm.year^−1^). The MATUR index decreased significantly with age in the entire cortex, except for the anterior part of the right insula, with the largest effects of age occurring in the frontal, parietal, temporal, and cingulate areas. This strong linear correlation with age was predominant in the right hemisphere, although the significance of this predominance was not assessed in the present study. Third, for the vertexwise quadratic age effect size, we found a three-to six-fold larger value in MATUR (from +0.36 to +1.76 μm.year^−2^) than in CT (from +0,12 to +0.27 μm.year^−2^). The network of areas showing the most substantial decrease in the maturation index in the youngest adults (evidenced by the quadratic modeling of age effect) covered the lateral cortex of the frontal lobes and the cingulate gyrus as well as the parietal areas. Using voxel-based morphometry (VBM, (Ashburner & Friston, 2000), Sowell et al. had previously studied the same brain regions during the maturational process from childhood to adolescence (Sowell et al., 2001) and to the beginning of adulthood (Sowell et al., 1999). These areas had the largest magnitudes of decrease in gray matter density. The effects of age on the present MATUR index indicate that frontal maturation occurs not only during adolescence but also during young adulthood and continues for many years. Moreover, the present results show that the regions with protracted maturation in the frontal cortex’s internal surface, from the motor areas to high-order associated areas supporting executive functions, exhibited the strongest negative correlation between CT and MYEL. Interestingly, in a population of young adults included in the HCP (age range 26-35 years), Thiebaut de Schotten et al. showed that the most anterior frontal areas, containing smaller cell bodies, were the thickest and the least myelinated (as observed in the present work) and had the highest entropy, which was defined as the number of resting-state intrinsic connections a region had with other cortical brain regions (Thiebaut de Schotten et al., 2017). This observation can be interpreted within the framework of Huntenburg et al.’s study (Huntenburg et al., 2017), which demonstrated a weak association between functional connectivity and MYEL in associative areas. Frontal maturation initiates during childhood and continues during young adulthood until the third decade of life. The region of the frontal lobe extending to the cingulate gyrus is linked to the general development of executive function, including attentional control goal setting and cognitive flexibility (Bush, Luu, & Posner, 2000). Moreover, the slower maturation of the right frontal lobe, which corresponded to the lowest maturation index in the younger participants, may be associated with the earlier development of this hemisphere’s gyral complexity [13], according to Dubois et al. (Dubois et al., 2010). In contrast, the motor and sensory, auditory, and primary visual areas showed smaller age effects and quadratic age effects. The cortical changes that we observed in the frontal and inferior parietal lobes and more prominently in the right hemisphere than in the left hemisphere before the age of 30 may play a role in the executive attentional network’s maturational process and self-regulation during this period (Rubia et al., 2010) (Westlye et al., 2011).

## 5. CONCLUSION

In the present study, we defined a multimodal cortical maturation index that reflects both cortical thickness and intracortical myelination and investigated its changes with age in a cross-sectional cohort of young and mature adults. This new neuroimaging marker was more sensitive to age-related cortical structural changes during adulthood than was either cortical thickness or myelin content alone. As evidenced by the large quadratic component of the age effect, the proposed maturation index dramatically characterizes the regional cortical territories exhibiting late brain maturation before stabilization occurs in late adulthood. These territories, essentially located in high-order frontal areas, support behavioral and cognitive self-regulation functions. In future investigations, this neuroimaging marker of late cortical maturation could help assess the impact of lifestyle factors, such as early alcohol and drug consumption, on the brains of young adults. An important goal for future research will also be to characterize the present maturational index’s changes across all life stages, from childhood to senescence.

## 7. ACKNOWLEDGMENTS

The authors are deeply indebted to Gael Jobard, Marc Joliot, Emmanuel Mellet, Laurent Petit, and Laure Zago for their contributions to the design and data acquisition of the data from the BIL&GIN. There are no conflicts of interest to declare.

